# InstructPLM: Aligning Protein Language Models to Follow Protein Structure Instructions

**DOI:** 10.1101/2024.04.17.589642

**Authors:** Jiezhong Qiu, Junde Xu, Jie Hu, Hanqun Cao, Liya Hou, Zijun Gao, Xinyi Zhou, Anni Li, Xiujuan Li, Bin Cui, Fei Yang, Shuang Peng, Ning Sun, Fangyu Wang, Aimin Pan, Jie Tang, Jieping Ye, Junyang Lin, Jin Tang, Xingxu Huang, Pheng Ann Heng, Guangyong Chen

**Author notes:** These authors contributed equally to this work (Jiezhong Qiu, Junde Xu, Jie Hu, Hanqun Cao).

## Abstract

Large language models are renowned for their efficacy in capturing intricate patterns, including co-evolutionary relationships, and underlying protein languages. However, current methodologies often fall short in illustrating the emergence of genomic insertions, duplications, and insertion/deletions (indels), which account for approximately 14% of human pathogenic mutations. Given that structure dictates function, mutated proteins with similar structures are more likely to persist throughout biological evolution. Motivated by this, we leverage cross-modality alignment and instruct fine-tuning techniques inspired by large language models to align a generative protein language model with protein structure instructions. Specifically, we present a method for generating variable-length and diverse proteins to explore and simulate the complex evolution of life, thereby expanding the repertoire of options for protein engineering. Our proposed protein LM-based approach, InstructPLM, demonstrates significant performance enhancements both in silico and in vitro. On native protein backbones, it achieves a perplexity of 2.68 and a sequence recovery rate of 57.51, surpassing Protein-MPNN by 39.2% and 25.1%, respectively. Furthermore, we validate the efficacy of our model by redesigning PETase and L-MDH. For PETase, all fifteen designed variable-length PETase exhibit depolymerization activity, with eleven surpassing the activity levels of the wild type. Regarding L-MDH, an enzyme lacking an experimentally determined structure, InstructPLM is able to design functional enzymes with an AF2-predicted structure. Code and model weights of InstructPLM are publicly available^*^.

## 1 Introduction

Mutations, encompassing point mutations arising from single nucleotide changes, as well as deletions, duplications, and insertions of multiple bases, serve as the driving force of evolution, facilitating the rapid adaptation of life to continuously changing environmental conditions as the fundamental cause of genetic diversity. These genetic variations provide a rich repository of proteins for reference and selection in protein engineering. A quintessential challenge in protein engineering is protein sequence design, also known as protein inverse folding, which entails identifying amino acid sequences capable of folding into a specified protein backbone structure. Highly accurate protein sequence design enables the generation of more effective enzymes [1, 2], improved protein-based therapeutics [3], and engineered proteins for industrial purposes like biofuels production and environmental remediation [4, 5]. The recent emergence of deep learning-based methods, exemplified by ProteinMPNN [6] and ESM-IF [7], has significantly advanced the field of protein sequence design. These methods formalize protein sequence design as a *multimodal learning* problem and solve the problem by training models with an encoder-decoder pattern which translates protein backbone structure into corresponding sequences. The swift evolution of deep learning architectures, including graph neural networks [6], and Transformers [8], coupled with the availability of high-quality protein structure data from Protein Data Bank (PDB) and AlphaFoldDB, has catalyzed substantial advancements in this field. For example, myoglobin and tobacco etch virus (TEV) protease designed by Protein-MPNN have demonstrated superior expression, stability, and functionality [9]; The de novo binders designed by ProteinMPNN exhibit high affinity and specificity to targets [10].

Multimodal learning, while straightforward and effective, faces a significant hurdle due to the paucity of high-quality cross-modal data. For instance, in the domain of visual-language understanding, training models from scratch necessitates access to well-curated datasets of image-text pairs, such as those of image captioning and visual question answering. Similarly, in protein sequence design, the training of the aforementioned ProteinMPNN depends on protein assemblies experimentally determined by X-ray crystallography or cryoEM in Protein Data Bank (PDB). To tackle the issue of limited data, an emerging strategy in the field of large vision-language models is *cross-modality alignment* [11–15], which is widely speculated to have been integrated into GPT-4[16], although the technical details of GPT-4 have yet to be publicly disclosed. This technique leverages existing pre-trained unimodal models to facilitate multimodal understanding. Pre-trained unimodal models learn domain-specific representations through extensive training on large unimodal datasets, yet they typically lack the ability to understand and reason across modalities. Cross-modality alignment bridges this gap by carefully aligning a pre-trained visual model with a pre-trained language model, thereby enabling effective cross-modal generation, understanding, and reasoning. Despite the success of cross-modality alignment techniques in vision-language understanding, their potential in protein sequence design remains unexplored due to the inherent dissimilarities between protein science and visual-language tasks. Consequently, addressing this challenge necessitates the development of powerful unimodal models and alignment techniques specifically tailored to proteins.

Fortunately, protein language models (pLMs), such as ESM [17], ProtGPT [18], and ProGen [19, 20], have emerged as a pivotal innovation in bioinformatics and computational biology, particularly in modeling protein sequences. Drawing inspiration from the achievements of large-scale language models like GPT and BERT, pLMs are pre-trained on vast datasets of protein sequences, leading to a comprehensive understanding of the protein universe. This includes the ability to capture the distribution of observed evolutionary sequences, generate novel yet viable sequences, and predict protein fitness. Specifically, pLMs have demonstrated the capability to generate functional protein sequences according to certain conditions. For example, GPT-based pLMs such as ProGen and ProtGPT can generate proteins following homologous samples or control tags specifying protein properties; ESM-based pLMs [21–23] design desired protein sequences by applying or sampling from the pre-trained masked language model. However, unlike general language models which exhibit zero-shot generalization and the ability to understand user intent on a wide range of tasks through methods like instruction fine-tuning [24, 25] or reinforcement learning [26, 27], it still remains an open area of inquiry how pLMs can generate protein sequences following fine-grained and complex biological instructions and even simulate the evolution of life. As mentioned in “Seven technologies to watch in 2024” [28] regarding deep learning for protein design, “Sequence-based approaches can build on and adapt existing protein features to form new frameworks, but they’re less effective for the bespoke design of structural elements or features.”.

In this work, we demonstrate the successful adaptation of *cross-modality alignment* and *instruction fine-tuning* techniques, originally developed for large language models, to the domain of protein sequence design. This transferability underscores the broad applicability of these techniques to life science. Our proposed model, InstructPLM, employs a lightweight cross-attention layer to align a frozen protein backbone encoder with a frozen protein language model decoder, aiming to teach the protein language model to design sequences following protein structure instructions. This configuration enables us to harness the robust generalization capabilities of pLMs, while stimulating pLMs to follow residue-level protein structure instructions at the same time. InstructPLM outperforms existing sequence design techniques in terms of perplexity and sequence recovery, while incurring the training of only an additional 2% of parameters.

The overall model architecture of InstructPLM is composed of three primary components — a protein language model decoder, a protein backbone encoder, and a protein structure-sequence adapter, as depicted in Figure 1 and described as follows:

**Fig. 1.**
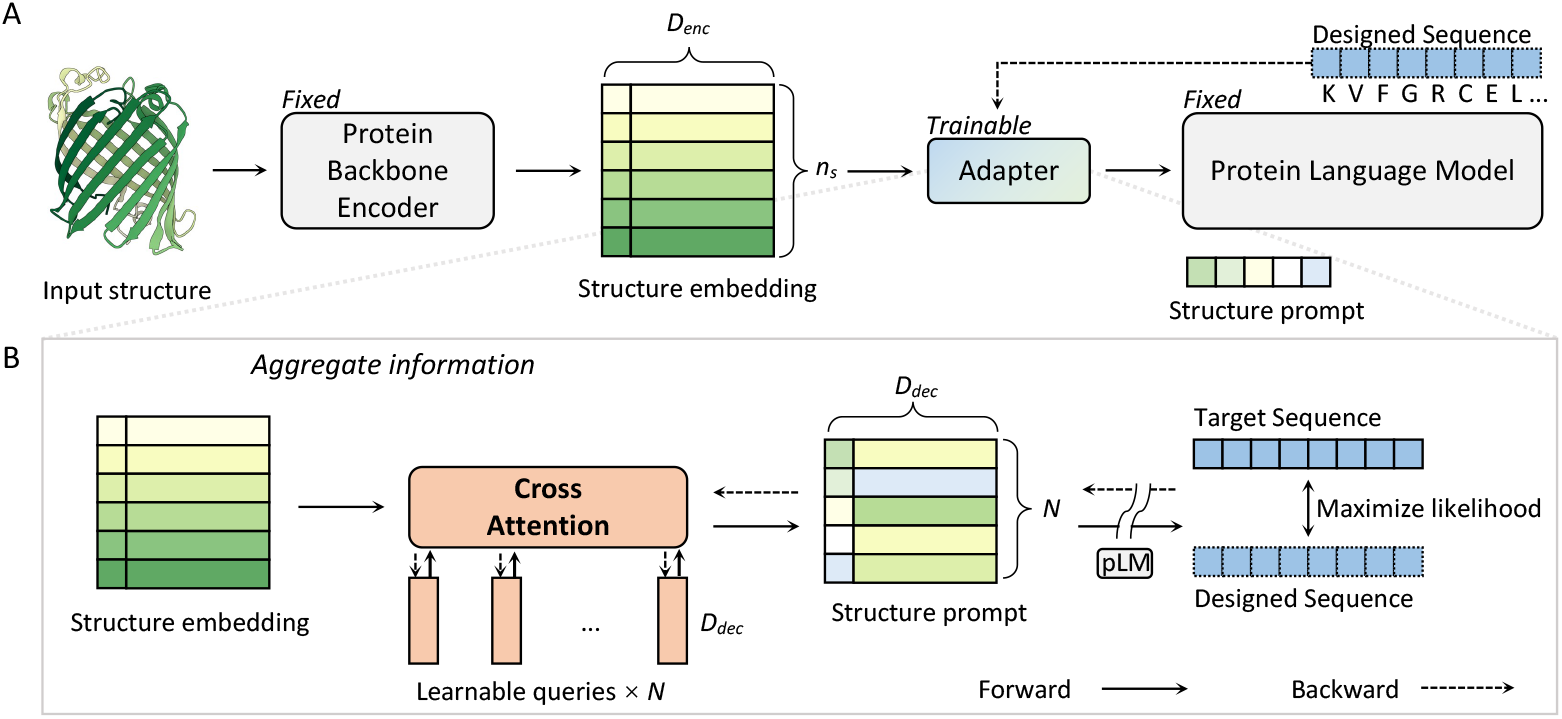
**(A)** Illustration of proposed method, InstructPLM. The network consists of three parts, the protein backbone encoder, the protein structure-sequence adapter, and the protein sequence decoder. The protein structure-sequence adapter is the only trainable part while the parameters of the structure encoder and sequence decoder are fixed during training. **(B)** Detailed illustration of the structure-sequence adapter. The adapter adopts a structure embedding as input, incorporating the structure information with a set of learnable queries. The queries will form as the structure prompt guiding the generation of the protein sequence decoder.

- **Protein Language Model Decoder**: InstructPLM adopts ProGen2 as its protein language model. The ProGen2 model family includes auto-regressive language models pre-trained on around 230 million protein sequences from a mixture of Uniref90 and BFD30 databases, whose model size ranges from 151M to 6.4B. We choose the largest one, ProGen2-xlarge with 6.4B parameters in InstructPLM. The ablation study about the size of the protein language model decoder is shown in Section 2.3.1.
- **Protein Backbone Encoder**: InstructPLM initializes its protein backbone encoder from other existing protein sequence design models, such as the backbone encoder of ProteinMPNN and ESM-IF. The ablation study about the choice of protein backbone encoder is shown in Section 2.3.2.
- **Protein Structure-Sequence Adapter**: The protein structure-sequence adapter is a key component of InstructPLM, not only because it is responsible for aligning structure and sequence into the same semantic space, but also because it contains all trainable parameters of InstructPLM. This adapter comprises a single-layer cross-attention module initialized randomly. The module uses several trainable embeddings as query vectors and the output of the protein backbone encoder as keys/values for cross-attention computation. This cross-attention module compresses the protein backbone embedding to a fixed-length structure instruction. The ablation about the number of queries (i.e., the length of structure instruction) is shown in Section 2.3.3. Additionally, we follow Qwen-VL [12] to add 1D absolute positional encodings into the cross-attention to keep protein primary structure information during compression. The compressed protein backbone structure feature sequence is subsequently fed into the protein language model as a soft prompt.

Our experimental results show that our InstructPLM framework is effective in protein sequence design in terms of perplexity and sequence recovery on the CATH dataset. Notably, we validate InstructPLM on redesigning PETase and L-MDH. For PETase, all fifteen designed variable-length proteins exhibit depolymerization activity, with twelve surpassing the activity levels of the wild type. For L-MDH, which does not have an experimentally determined structure, we show that InstructPLM is able to design functional enzyme with AlphaFold2-predicted structure — 3 out of 15 designed L-MDH exhibit detectable enzymatic activity.

## 2 Results

We firstly evaluate InstructPLM by following the standard in silico protein sequence design benchmark in Section 2.1. The experimental evaluation and ablation studies are then described in Section 2.2 and Section 2.3.

### 2.1 In Silico Protein Sequence Design

#### 2.1.1 InstructPLM Designs Sequences with High Recovery

For computational evaluation, we assess the performance of InstructPLM using CATH 4.2 [35], which is a widely recognized benchmark in protein sequence design. We adopt the official train, validation, and test splits, which contain 18,024, 608, and 1,120 proteins respectively. Table 1 and the left panel of Figure 2(A) show the detailed sequence design performance on the CATH 4.2 held-out test split.

**Table 1.**
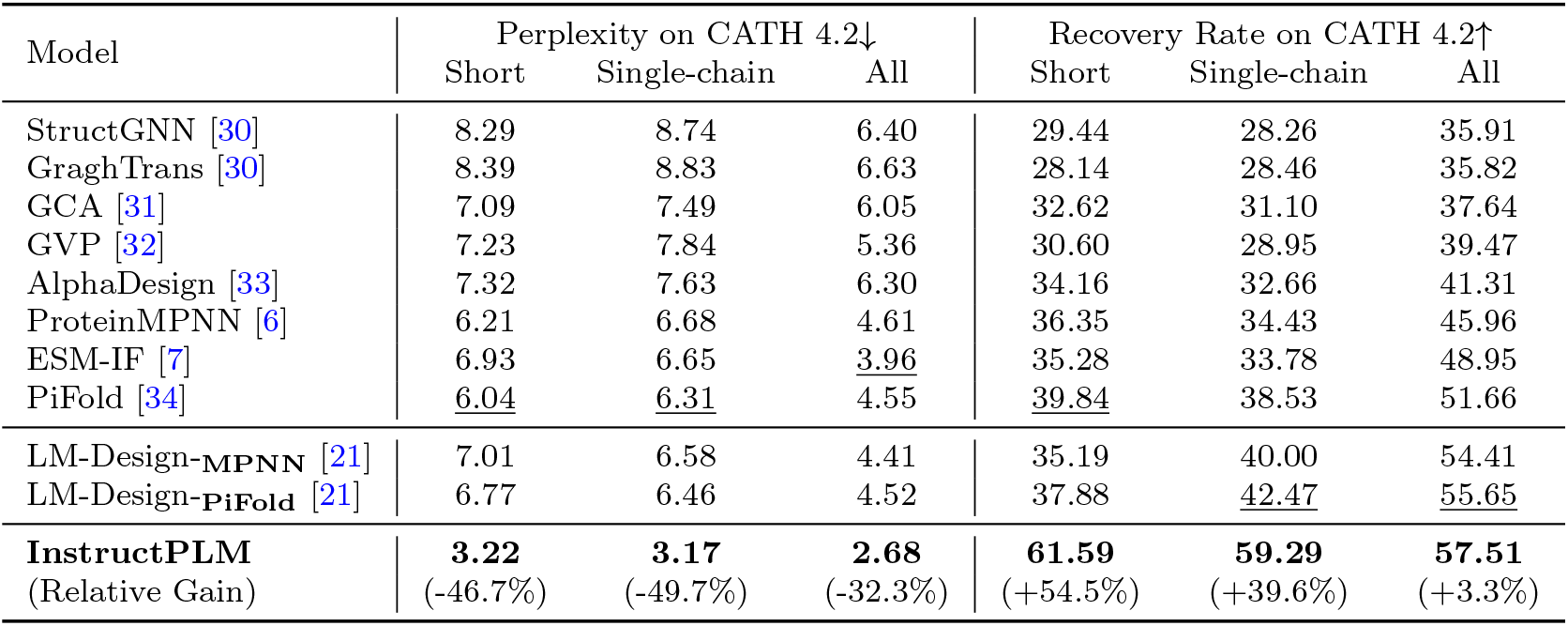
Sequence design performance on CATH 4.2 held-out test split. The metrics include perplexity (exponentiated categorical cross-entropy loss per residue) and recovery rate (percentage of correctly predicted residues). The best performance is shown in **bold**, while the best baseline is indicated with an underline. For a fair comparison, we test the performance of ESM-IF on CATH 4.2, although it is originally trained and evaluated on CATH 4.3.

| Model | Perplexity on CATH 4.2↓ |  |  | Recovery Rate on CATH 4.2↑ |  |  |
| --- | --- | --- | --- | --- | --- | --- |
|  | Short | Single-chain | All | Short | Single-chain | All |
| StructGNN [30] | 8.29 | 8.74 | 6.40 | 29.44 | 28.26 | 35.91 |
| GraghTrans [30] | 8.39 | 8.83 | 6.63 | 28.14 | 28.46 | 35.82 |
| GCA [31] | 7.09 | 7.49 | 6.05 | 32.62 | 31.10 | 37.64 |
| GVP [32] | 7.23 | 7.84 | 5.36 | 30.60 | 28.95 | 39.47 |
| AlphaDesign [33] | 7.32 | 7.63 | 6.30 | 34.16 | 32.66 | 41.31 |
| ProteinMPNN [6] | 6.21 | 6.68 | 4.61 | 36.35 | 34.43 | 45.96 |
| ESM-IF [7] | 6.93 | 6.65 | <u>3.96</u> | 35.28 | 33.78 | 48.95 |
| PiFold [34] | <u>6.04</u> | <u>6.31</u> | 4.55 | <u>39.84</u> | 38.53 | 51.66 |
| LM-Design-MPNN [21] | 7.01 | 6.58 | 4.41 | 35.19 | 40.00 | 54.41 |
| LM-Design-PiFold [21] | 6.77 | 6.46 | 4.52 | 37.88 | <u>42.47</u> | <u>55.65</u> |
| **InstructPLM**<br>(Relative Gain) | **3.22**<br>(-46.7%) | **3.17**<br>(-49.7%) | **2.68**<br>(-32.3%) | **61.59**<br>(+54.5%) | **59.29**<br>(+39.6%) | **57.51**<br>(+3.3%) |

**Fig. 2.**
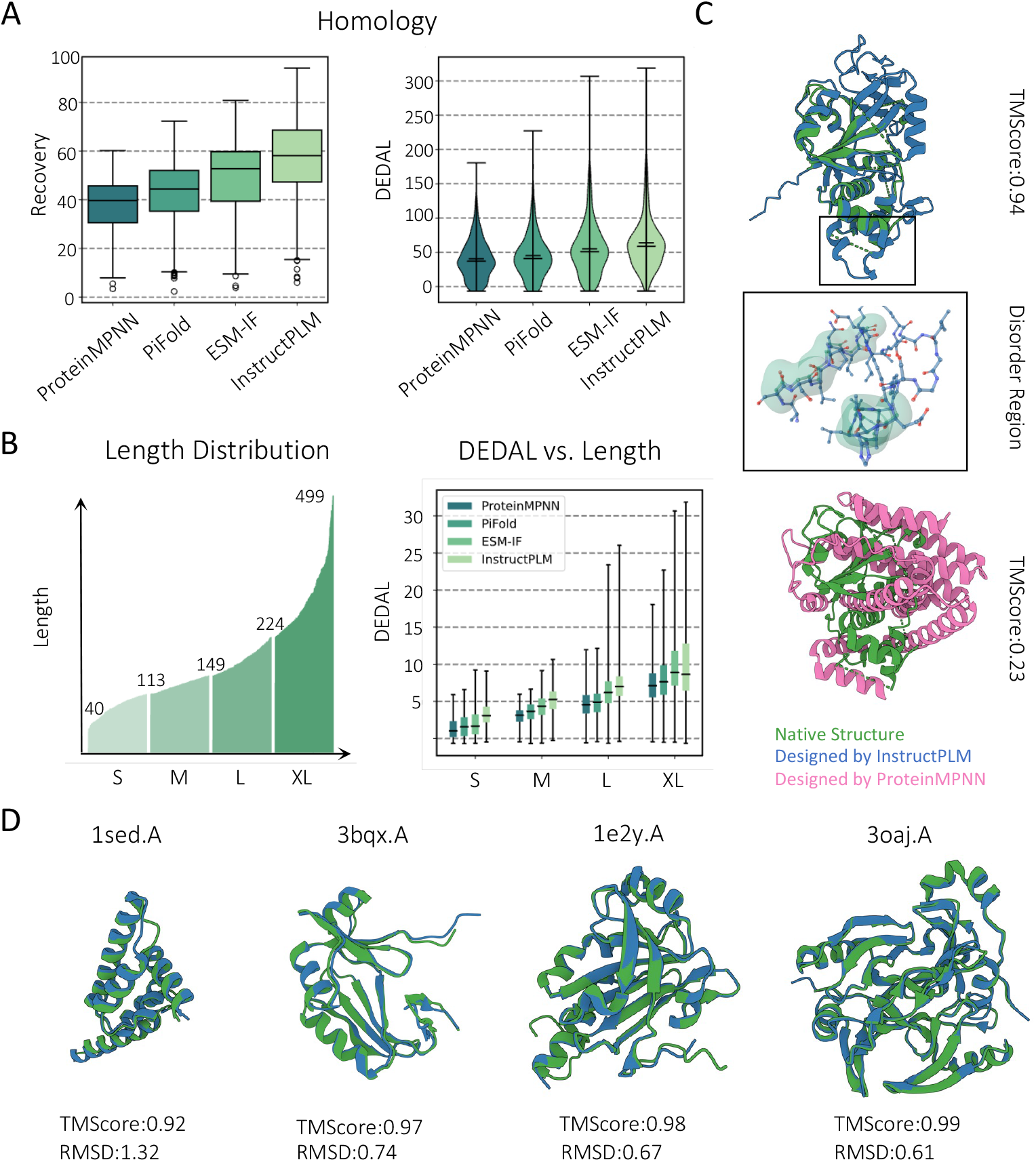
In silico protein sequence design performance of InstructPLM. **(A)** Recovery rate (left) and homology score predicted by DEDAL (right) on CATH 4.2 test split. Compared to existing methods, InstructPLM achieves the best performance. **(B)** DEDAL score breakdown on different sequence lengths. InstructPLM achieves consistently better performance than other approaches. **(C)** Comparison of predicted structures of the designed sequence of PDB code 4F0D. Compared to ProteinMPNN (TM-Score: 0.23), InstructPLM (TM-Score: 0.94) generates more plausible sequences in the regions where the protein structure is missing. **(D)** Additional designed cases from InstructPLM, using CATH 4.2 test set as the structure templates. All TM-Scores are computed using TM-align [29].

InstructPLM demonstrates superior performance compared to models trained from scratch on the CATH 4.2 dataset. Regarding perplexity, InstructPLM significantly improves upon the best baseline method, ESM-IF, by 32.3% (from 3.96 to 2.68). In sequence recovery, InstructPLM also shows a significant enhancement, with an 11.3% increase in the recovery rate over the best baseline, PiFold (from 51.66 to 57.51). It’s particularly noteworthy that the comparison between InstructPLM and ProteinMPNN highlights the efficacy of protein language models in representing the distribution of evolutionary sequences. This is evident as they utilize the same backbone encoder, with the primary distinction lying in their sequence decoders.

The recently introduced LM-Design [21] shares a conceptual connection with InstructPLM, as both leverage protein language models (pLMs) for the generation of protein sequences. The distinction between them primarily hinges on their approach to decoding protein sequences. LM-Design employs a modified ESM pLM in a step-wise fashion to iteratively refine a sequence initially decoded by ProteinMPNN or PiFold, utilizing the masked language model of ESM. On the other hand, InstructPLM decodes protein sequences in an autoregressive manner, utilizing the protein backbone embedding derived from ProteinMPNN. A comparative analysis of InstructPLM and LM-Design reveals the benefits of the cross-modality alignment and instruction tuning techniques that we have incorporated. These techniques enable a comprehensive exploitation and effective use of structural information in sequence design. Specifically, when comparing InstructPLM with LM-Design-_**MPNN**_, which both leverage Protein-MPNN, InstructPLM demonstrates relative improvements of 39.2% (from 4.41 to 2.68) in perplexity and 5.7% (from 54.41 to 57.51) in recovery rate, respectively.

More interestingly, InstructPLM exhibits a consistent pattern of improvement across different subsets of CATH 4.2, and especially achieves more significant enhancements when designing short proteins and single-chain proteins. This observation suggests that the model’s architecture, training techniques, and the pLM’s prior knowledge are particularly effective for these protein types. This also indicates that InstructPLM is not only a powerful general-purpose protein designer but also excels in specialized applications, particularly where the protein sequences are shorter or consist of a single chain.

We also evaluate InstructPLM on TS50 and TS500 datasets, which consist of 50 and 470 proteins and are often employed as additional benchmarks to further test generalization capability [21, 33, 34] beyond CATH dataset. The detailed results are shown in Table 7 in Appendix, where InstructPLM demonstrates consistent and robust performance.

#### 2.1.2 InstructPLM Generates Variable-Length and Homologous Sequences

In contrast to conventional approaches that generate sequences of a fixed length, a distinctive feature of InstructPLM lies in its ability to create protein sequences of variable lengths through an open-ended generation process. This is facilitated by the aligned pLM decoder. It generates subsequent amino acid tokens in an autoregressive manner, pausing only upon encountering the termination token. This innovative capacity for open-ended generation aligns closely with the intrinsic nature of biological evolution, where a variety of mutations such as insertions, deletions, and their combinations are commonplace. However, the traditional sequence recovery measure is not well-suited for evaluating the quality of variable-length sequences, as it evaluates the per-position accuracy of designed sequences compared to their native counterparts. To effectively assess InstructPLM’s unique capability, we propose to use sequence homology score as a new metric. We employ DEDAL [36], a state-of-the-art deep learning-based method for homology detection. This method has shown a remarkable improvement in alignment accuracy, achieving up to two or three times better performance than traditional methods when searching for remote homologs.

Within the test split of CATH 4.2, we randomly design 20 protein sequences at a temperature setting of 0.15 for each native protein structure. The sequence homology score, as determined by DEDAL, serves as a metric for comparing the similarity between the native and artificially designed sequences. As depicted in the right panel of Figure 2(A), the distribution of homology scores across various methods is showcased, with InstructPLM achieving an average homology score of 63.97, surpassing ProteinMPNN/ESM-IF/PiFold by margins of 23.25/8.83/18.9, respectively. Figure 2(B) provides a detailed breakdown of homology score distributions across sequences of different lengths, demonstrating that InstructPLM maintains a consistently high level of performance in the design of sequences with varying lengths.

### 2.2 Experimental Evaluation of InstructPLM

Although InstructPLM achieves significant improvement regarding in silico sequence design metrics such as perplexity, sequence recovery, and homology score, we aim to test its ability to design functional proteins. Specifically, we evaluate InstructPLM on PETase (PDB code: 7SH6) and L-MDH (Uniprot: A0A319AA41). Notably, L-MDH has no experimentally determined structure, thus we design L-MDH based on its AF2-predicted structure. The design process is outlined in Figure 3(A). Utilizing the backbone structure as a starting point, InstructPLM produces 10,000 potential sequences via auto-regressive generation with top-p sampling (*p* = 0.9) and a temperature of 0.8, as detailed in Section 3.3. The sequences’ structures are predicted using ESMFold [37] and compared to the native structure with DeepScore from DeepAlign package [38, 39] to calculate TM-Score, which is a measure of structural similarity. Lastly, we choose the top sequences ranked by TM-Score for further experimentation. Upon meticulous experimental evaluation, the sequences designed by our model for the two proteins showed activity, demonstrating the model’s effectiveness. We provide a detailed study of each designed enzyme in Section 2.2.1 and Section 2.2.2

**Fig. 3.**
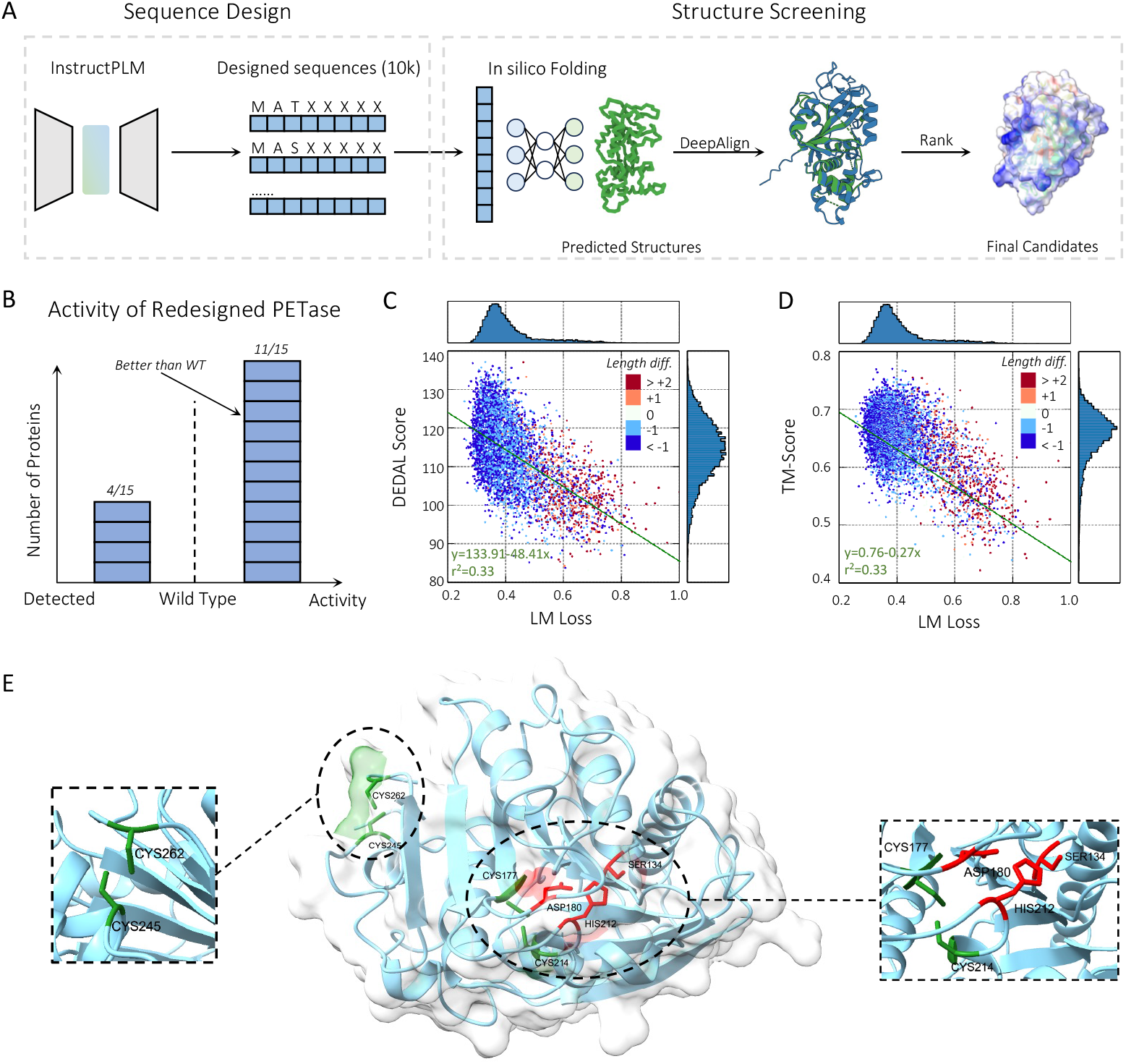
Experimental evaluation of InstructPLM. **(A)** Overview of the protein design post-processing pipeline using InstructPLM. Given a target structure, InstructPLM samples 10,000 candidate sequences. The structures of all sequences are predicted using ESMFold. DeepAlign is then employed for structure alignment. Finally, samples with higher structural similarity are selected for further experiments. **(B)** Result of PETase. All 15 sequences designed by InstructPLM exhibit high expression levels. **(C)** The joint distribution of the homology score (DEDAL score) and language model loss (LM loss) indicating the overall design quality of the 10,000 candidate sequences. **(D)** The joint distribution of the structure similarity score (TM-Score) and language model loss (LM loss) providing an overview of the 10,000 designed sequences. **(E)** An visualization case of PETase designed by InstructPLM, the protein structure is predicted by ESMFold. It has catalytic triad SER134-HIS212-ASP180 and two disulfide bridges (DS1 and DS2).

**Fig. 4.**
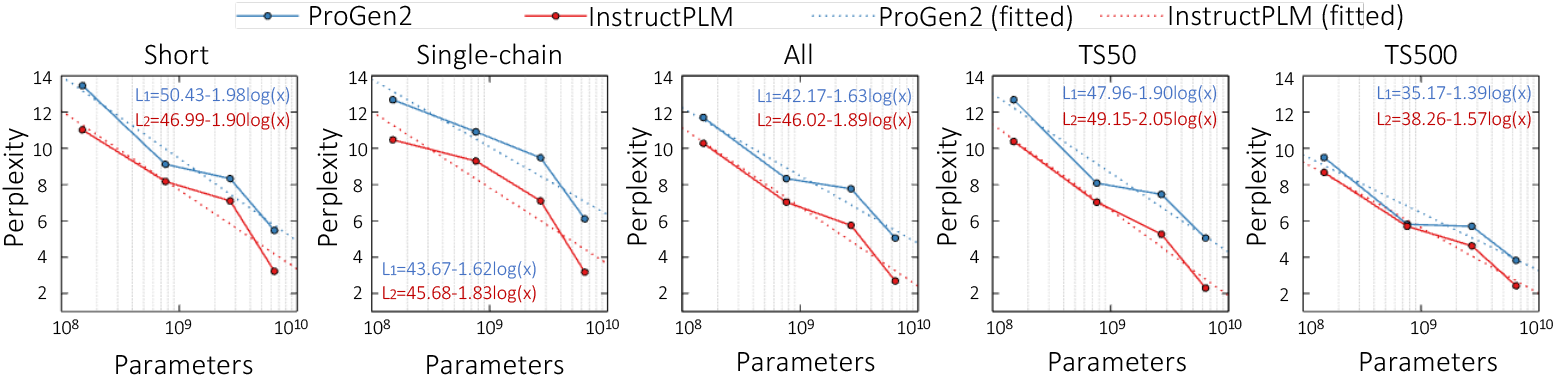
Scaling law of InstructPLM w.r.t. perplexity on CATH 4.2 and TS50/500 respectively.

**Fig. 5.**
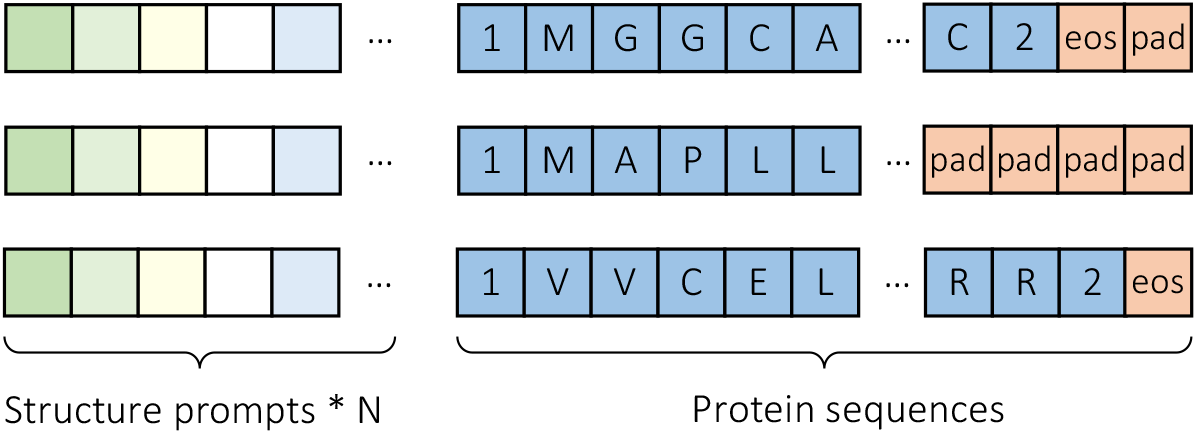
Illustration of batch construction. We concat the structure prompt with protein sequence to train the InstructPLM, and special tokens ‘1’, ‘2’, ⟨*eos*⟩ and ⟨*pad*⟩.

#### 2.2.1 PETase

Designing PETase poses a test of InstructPLM’s ability to improve upon naturally occurring proteins. We select the top 15 sequences in terms of TM-Scores for experimentation. The lengths of these 15 sequences vary from 261 to 269, reflecting the InstructPLM’s capability to generate sequences of variable lengths.

When expressed in Escherichia coli, all 15 sequences designed by InstructPLM show good expression levels. The PET hydrolysis reaction products — bis (2-hydroxyethyl) terephthalate (BHET), mono (2-hydroxyethyl) terephthalate (MHET), and terephthalic acid (TPA) — are quantified using high-performance liquid chromatography. All 15 sequences designed by InstructPLM have PET hydrolytic activity, while 11 sequences are better than wild-type (PDB code: 5XJH), as shown in Figure 3(B). Sequence alignment, as shown in Figure 6, reveals that all 15 designed PETase sequences contain the strictly conserved catalytic triad SER134-HIS212-ASP180. SER134 serves as the nucleophile and is located within hydrogen-bond distance to be polarized by the base HIS212, which is in turn stabilized by acid ASP180 [40]. Additionally, 4 out of 15 sequences have two intramolecular disulfide bridges (DS1:C177-C214 and DS2:C245-C262), and all (15/15) have DS2 bridge connecting the C-terminal helix at the last loop. An example of a sequence generated by InstructPLM featuring both the catalytic triad and the two disulfide bridges is shown in Figure 3(E).

Through this comprehensive analysis, we observe that InstructPLM has effectively learned the significance of catalytic and disulfide bridge residues, which is crucial for enhancing the success rate of designed PETase enzymes. To further our understanding, we perform two regression analyses to explore the correlation between (1) language model loss (LM loss) and homology score (DEDAL score); and (2) LM loss and TM-Score. The results, depicted in Figures 3(C) and 3(D), suggest that LM loss can be a valuable metric for assessing the similarity in both sequence and structure between designed and native proteins.

#### 2.2.2 L-MDH

L-MDH (L-Malate dehydrogenase, Uniprot: A0A319AA41) is a key enzyme in the citric acid cycle with no experimentally determined structure available. We rely on AlphaFold2 [41] predicted structures as the basis for design. Following the same protocol of PETase, we generate 10,000 sequences with InstructPLM, and select 15 sequences with the highest TM-Scores. Out of the 15 designed L-MDH sequences, our experimental validation reveals that three exhibited detectable enzymatic activity. While these active variants do not surpass the performance of the wild-type enzyme, the fact that they demonstrated activity is a significant achievement, given the absence of an experimentally confirmed structure for L-MDH. The ability to generate functional enzymes based solely on computationally predicted structures underscores the potential of combining advanced protein structure prediction tools like AlphaFold [41] and structure design tools such as Chroma [42] with our design methodology. Notably, both three sequences are shorter than wild-type, further demonstrating the flexibility and adaptability of our approach. We compare the L-MDH with three detectable enzymes by sequence alignment and found that all of them have the same activate site HIS179 which hit the UniRule PIRSR: PIRSR000102-1 [43] (See Figure 8). An example of L-MDH structure predicted by ESMFold [37] is shown in Figure 7.

**Fig. 6.**
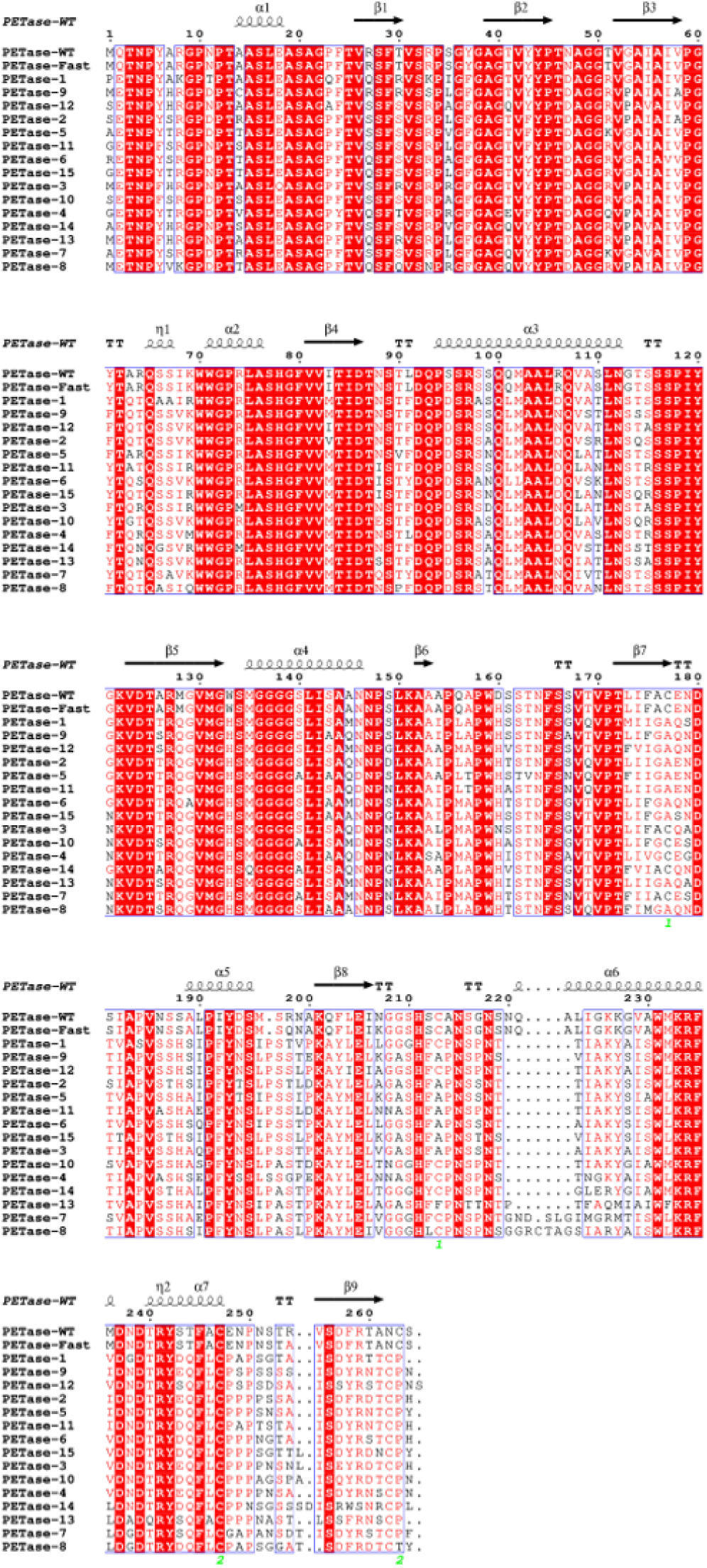
Alignment by ENDscript [52] of PETase generated by InstructPLM, Fast-PETase[53] and Wiled-type PETase. All of them (15/15) have the strictly conserved catalytic triad SER134-HIS212-ASP180. Not all sequences (4/15) have two intramolecular disulfide bridges (DS1 and DS2) and all of them (15/15) have DS2 connects the C-terminal helix and the last loop.

**Fig. 7.**
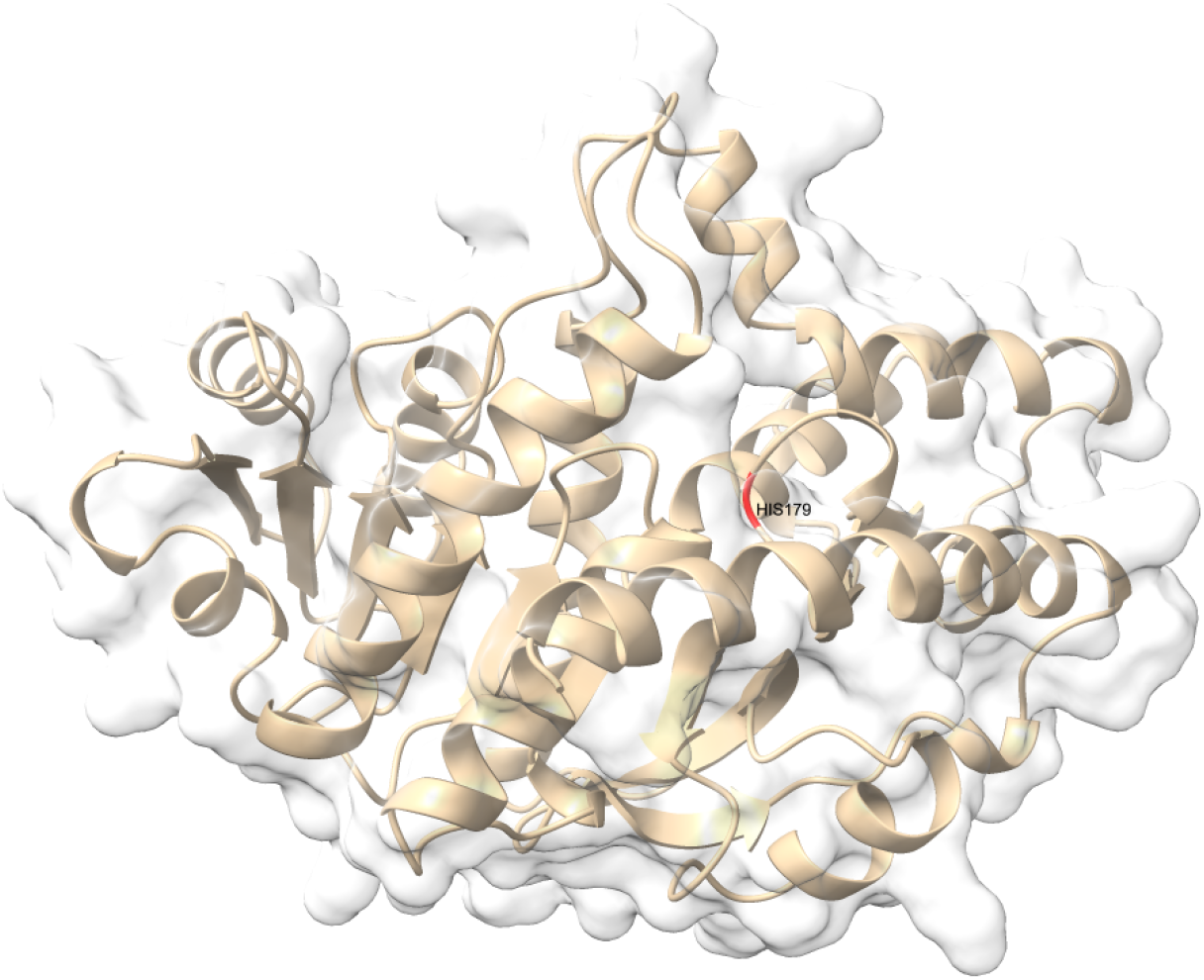
An example of L-MDH designed by InstructPLM, the protein structure is predicted by ESMFold. It has the same activate site HIS179 with Uniprot: A0A319AA41 which hit the UniRule PIRSR: PIRSR000102-1 [54].

**Fig. 8.**
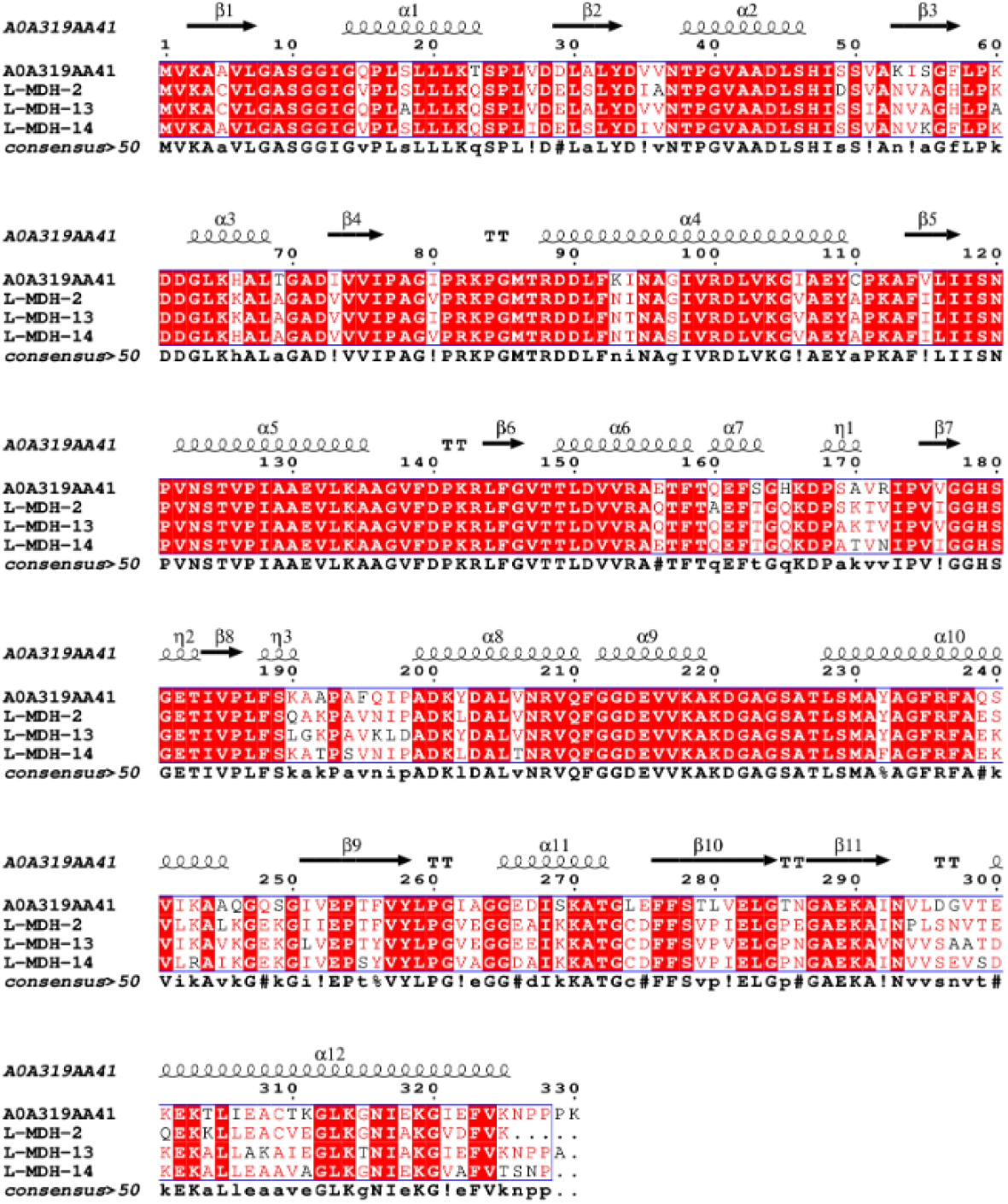
Alignment by ENDscript [52] of L-MDH(Uniprot:A0A319AA41) and L-MDH which generate by InstructPLM and have detectable enzymatic activity. It has the same activate site HIS179 with Uniprot: A0A319AA41 which hit the UniRule PIRSR: PIRSR000102-1 [54].

### 2.3 Ablation Studies and Discussion on Model Designs

#### 2.3.1 Scaling Law of InstructPLM

Scaling laws [45], in the context of large language models, refer to the observation that the next token prediction loss of these models scales as a power-law in terms of the number of parameters, the amount of training data, or the computational resources used during training. In this work, we mainly investigate the relationship between the perplexity of InstructPLM and the model size of its ProGen2 pLM decoder. The ProGen2 model family [44] includes autoregressive pLMs whose model size ranges from 151M to 6.4B. We conduct an ablation study by enumerating pLM decoder in InstructPLM from ProGen2-small (151M parameters), ProGen2-base (764M parameters), ProGen2-large (2.7B parameters), and ProGen2-xlarge (6.4B parameters). All other model configurations are the same as Section 3.2. The experimental results are shown in Table 2 and Figure 4. On all five datasets, both ProGen and InstructPLM follow the scaling law of large language models — their language model loss can be predicted using a power-law with respect to the model size. Compared to ProGen2, InstructPLM has achieved stable and consistent improvements across different model sizes and various datasets. This not only demonstrates the effectiveness of cross-modality alignment and instruction fine-tuning techniques we adopted in InstructPLM but also suggests that protein sequence design tasks can continuously benefit from the scaling of protein language models towards better performance.

**Table 2.** Ablation results on pre-trained pLMs: the perplexity of CATH 4.2, TS50, and TS500 is obtained based on different pre-trained ProGen2 models [44]. The number of trainable parameters and the total parameters are shown. The best performance is shown in **bold**.

| Method | CATH 4.2 (test split) |  |  | TS50 | TS500 | Trainable/Total |
| --- | --- | --- | --- | --- | --- | --- |
|  | Short | Single-chain | All |  |  |  |
| ProGen2-small [44] | 13.46 | 12.68 | 11.70 | 12.68 | 9.49 | - |
| InstructPLM-small | 11.02 | 10.47 | 10.28 | 10.38 | 8.67 | 5.9M/306M(2.0%) |
| ProGen2-base [44] | 9.12 | 10.91 | 8.33 | 8.08 | 5.81 | - |
| InstructPLM-base | 8.17 | 9.30 | 7.03 | 7.03 | 5.70 | 13.7M/927M(1.5%) |
| ProGen2-large [44] | 8.33 | 9.48 | 7.77 | 7.46 | 5.70 | - |
| InstructPLM-large | 7.10 | 7.10 | 5.75 | 5.26 | 4.62 | 34.6M/2.9B(1.2%) |
| ProGen2-xlarge [44] | 5.47 | 6.11 | 5.05 | 5.05 | 3.82 | - |
| InstructPLM-xlarge | **3.22** | **3.17** | **2.68** | **2.29** | **2.42** | 89.1M/6.6B(1.3%) |

#### 2.3.2 Ablating Protein Backbone Encoder

We further ablate the usage of different protein backbone encoders in InstructPLM, including the encoders of ProteinMPNN [6], ESM-IF [7] and PiFold [34]. For ProteinMPNN, we concatenate the protein backbone representations encoded by nine released ProteinMPNN models^1^, each with a dimension of 128, and finally obtain protein backbone embedding with a dimension of 128×9 = 1152. For ESM-IF and PiFold, we adopt their backbone encoders which represent protein backbone structures to embeddings with dimensions of 512 and 128, respectively. Notably, although ESM-IF is originally trained on CATH 4.3, we present results of InstructPLM-_**ESM**−**IF**_ trained on the CATH 4.2 dataset, mainly for a fair comparison. The experimental results are detailed in Table 3. Among all datasets, we observe that InstructPLM-_**MPNN**_ yields the best performance, showing the robustness and effectiveness of ProteinMPNN.

**Table 3.** Ablation study on protein backbone encoder: The perplexity of InstructPLM with different protein backbone encoders and different types of embeddings on CATH 4.2 test split, TS50, and TS500. The best performance is shown in **bold**.

| Method | Dimension of Backbone Emb. | CATH 4.2 (test split) |  |  | TS50 | TS500 |
| --- | --- | --- | --- | --- | --- | --- |
|  |  | Short | Single-chain | All |  |  |
| InstructPLM-**PiFold** | 128 | 3.70 | 3.81 | 3.28 | 4.62 | 3.74 |
| InstructPLM-**ESM-IF** | 512 | 4.44 | 4.34 | 3.24 | 2.88 | 2.71 |
| InstructPLM-**MPNN** | 1152 | **3.22** | **3.17** | **2.68** | **2.29** | **2.42** |

#### 2.3.3 Ablating the Number of Learnable Queries in Adapter

To probe the expression power of the protein structure-sequence adapter for sequence design, we conduct an ablation study on the number of learnable queries in the cross-attention of the adapter. We demonstrate the sequence design performance in terms of perplexity on the test split of CATH 4.2, TS50, and TS500 datasets. With other hyper-parameters the same as the training phase, the query length is chosen from 32, 64, 128, 256, and 512 respectively. The detailed performance of InstructPLM with different numbers of queries is shown in Table 4. InstructPLM achieves better perplexity as the increase of query length, illustrating how protein structure-sequence adapter compresses and provides meaningful instructions to the ProGen2 decoder. However, increasing the number of queries beyond 256 may lead to over-fitting. Although InstructPLM with 512 queries achieves better perplexity than InstructPLM with 256 queries on two subsets of CATH 4.2 (i.e., Short and Single-chain), we can observe significant performance drops on CATH 4.2 (All), TS50 and TS500. To utilize more robust models, we set the number of queries to 256 by default.

**Table 4.**
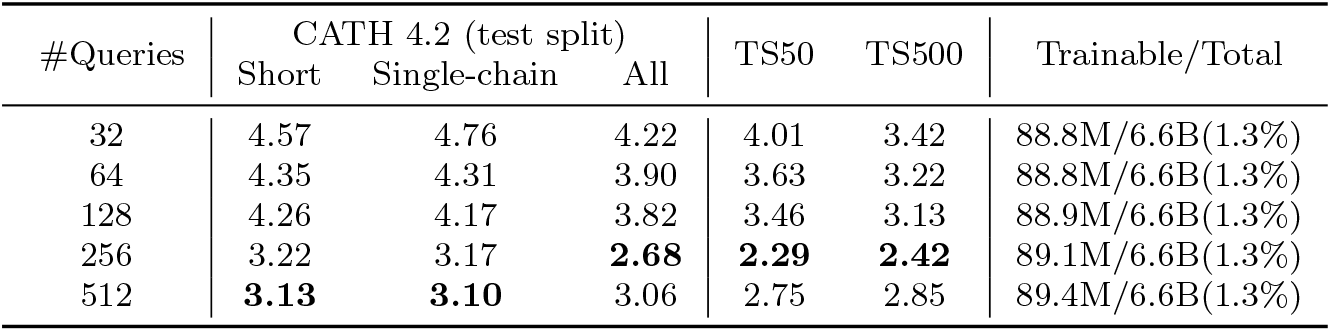
Ablation results on structure query length: the perplexity of CATH 4.2, TS50, and TS500 is obtained based on structure guidance of ProteinMPNN [6] embeddings. The amount of trainable parameters and total parameters are shown. The best performance is shown in **bold**.

## 3 Method

### 3.1 Model Architecture

For an input protein backbone, we aim to identify a suitable protein sequence *S* where *S* = (*s*_1_, *s*_2_, …) which folds to the given structure. The overall framework of InstructPLM contains three main components, including the pre-trained protein backbone encoder, the protein structure-sequence adapter, and the pre-trained protein language model decoder. We present our models step by step.

#### 3.1.1 Protein Backbone Encoder

We first obtain structure embedding 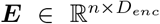 with a pre-trained structure encoder, where *n* is the number of protein residues and *D*_*enc*_ is the embedding dimension of the structure encoder. The embedding dimension *D*_*enc*_ can vary for different encoders. We utilize structure encoders from well-established protein sequence design methods, such as ProteinMPNN [6], ESM-IF [7], and PiFold [34]. To initialize these models, we make use of the officially released pre-trained weights. Specifically, for ProteinMPNN [6], we concatenate the embeddings from its nine models, which include four vanilla models, two soluble models, and three *C*_*α*_ models.

#### 3.1.2 Protein Structure-Sequence Adapter

The protein structure-sequence adapter consists of four main components: linear projection layers for aligning structural embeddings to a latent interaction space, a dynamic positional encoding layer to capture residue-level connections for proteins, a cross-attention layer with a set of learnable queries 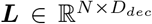 for knowledge transfer, where *N* is a tunable hyper-parameter and *D*_*dec*_ is determined by hidden dimension of pre-trained sequence decoder. Finally, we deploy an output projection layer that maps to language embeddings. To ensure the stability of module outputs, we further incorporate three normalization layers.

For an input structure embedding ***E***, the adapter will pad ***E*** with zeros to a fixed length *n*_*s*_. We set *n*_*s*_ to 512 by default. We first incorporate positional information into the structure embedding ***E*** and learnable queries ***L***. For protein sequences with max length of *n*_*s*_, the positional encoding ProtPE is defined for each dimension *i* = 0, …, *D*_*dec*_*/*2 − 1 of the encoding at position *pos* = 0, … *n*_*s*_ − 1 as:

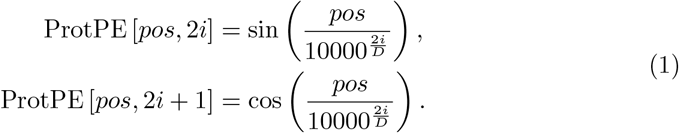

The structure embedding ***E*** is projected to the latent interaction space before coupling with positional embedding:

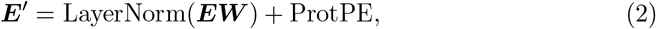

Where 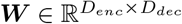. As the length of learnable queries ***L*** is inconsistent with the input structure length *n*_*s*_, the positional information is injected into the queries by adding the positional feature embeddings with evenly distributed stride:

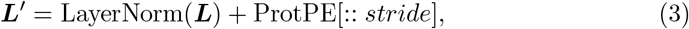

where the *stride* = *n*_*s*_*//N* .

The cross-attention layer is formulated as a function that maps queries, keys, and values to outputs. We define keys and values according to the structural embeddings in Equation 2 and query them with the learnable query as defined i n E quation 3. To enhance the model’s capacity to attend to information from different sub-spaces, we employ a multi-head attention strategy [8], where the attention function is performed in parallel across *A* different attention heads. For e ach a ttention head *k* ∈ {1, 2, …}, *A*, the structural embeddings and queries are projected into the same dimension by linear projections:

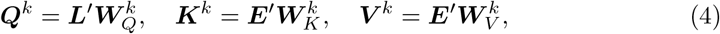

where 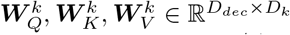 are parameter matrices for queries, keys, and values, respectively, and *D*_*k*_ = *D*_*dec*_*/A* represents the dimension of attention heads.

To transform the information from structural embeddings into the pre-defined queries, we obtain the interaction score based on inner products:

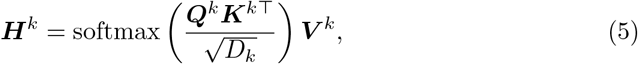

This operation computes the similarity between the queries and keys, which is then used to weight the values, effectively transferring the structural information to the outputs. The multi-head attention outputs are then concatenated across heads and linearly transformed to the final outputs of cross-attention:

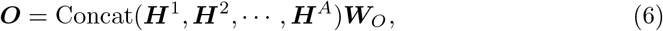

where each ***H***^*k*^ denotes independent outputs from a specific attention head, and 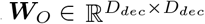 is the parameter matrix for the final linear transformation.

Finally, a normalization layer and a linear projection layer 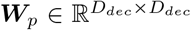 are adopted to transform the attention output to the prompt ***P*** of the pLMs,

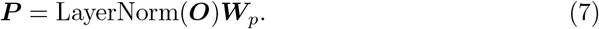

Further details regarding the model’s weight initialization scheme and training hyperparameter settings are shown in the Appendix 6.

#### 3.1.3 Protein Language Model Decoder

We utilize a pre-trained large protein language model as our sequence decoder. Specifically, we use ProGen2 [44] as our decoder, as ProGen2 is currently the largest and open-sourced auto-regressive protein language model. Given the structure prompt ***P*** obtained by previous steps, we concatenate the prompt ***P*** with the special token ‘1’ (following ProGen2, where ‘1’ indicates the N-terminal side of the sequence.) and send it to the sequence decoder model. The decoder generates the protein sequence from the conditional probability distribution of the next token in an auto-regressive manner,

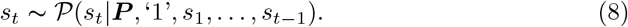

For the fixed-length generation, we auto-regressively sample from the probability distribution in Equation 8 for *n* iterations. While for open-ended generation, we auto-regressively sample from the probability distribution in Equation 8 until encountering the C-terminal token, namely ‘2’. By integrating these components and techniques, InstructPLM efficiently processes protein structural embeddings on high-dimensional latent space and generate corresponding protein sequences.

### 3.2 Training

The InstructPLM is trained on the CATH 4.2 training set and validated on the CATH 4.2 validation set, no other data are included during training, suggesting the data efficiency of our methods. The training objective is to minimize the cross-entropy loss at the protein-token level, which measures the discrepancy between the predicted and ground-truth amino acid sequences.

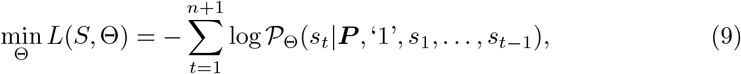

where Θ represents the trainable parameters of the model, all from the protein structure-sequence adapter. The logarithm term computes the log-likelihood of the amino acid *s*_*t*_ given the protein structure prompt ***P*** and the preceding sequence tokens ‘1’, *s*_1_, …, *s*_*t*−1_. In addition, we also minimize the loss of the special C-terminal side token ‘2’ to ensure InstructPLM can stop properly at the end of sequences. Thus we append the special token ‘2’ at the end, i.e., *s*_*n*+1_ = ‘2’.

We use Adam [46] optimizer with a maximum learning rate of 1*e*^−4^, and a constant learning rate warm-up is applied for the first 5,000 steps to stabilize the learning process. The training procedure is performed on four NVIDIA A100 GPUs, with a batch size of 16 on each GPU, and using BF16 precision, enabling efficient parallel processing and reduced training time. The whole training configuration is summarized in Table 5. By leveraging this training methodology, InstructPLM learns to effectively capture the relationship between protein structures and sequences, enabling accurate generation of sequences from given structural information. The frozen pre-trained ProGen2 provides a solid foundation, while the adapter focuses on learning the cross-modality alignment, offering a powerful solution to protein sequence design.

**Table 5.** Summary of model hyperparameters.

| Hyperparameter | Value | Description |
| --- | --- | --- |
| *Model* |  |  |
| Max length | 512 | Max number of amino acids of input structures. |
| Learnable Queires | 256 | Number of learnable queries. |
| Number of Heads | 16 | Number of heads in multi-head cross attention. |
| *Generation* |  |  |
| Temperature | 0.15/0.8 | Control the diversity of generated sequences. |
| Top-p | 0.9 | Control the diversity of generated sequences. |
| *Training* |  |  |
| Learning Rate | 0.0001 | Learning rate for the optimizer. |
| Batch Size | 16 | Number of samples per step. |
| Data Parallel | 4 | Number of batches per step. |
| Epochs | 200 | Number of times to iterate over the entire data-set. |
| Warmup Steps | 5000 | Number of steps to gradually increase the learning rate at the beginning of training. |
| Weight Decay | 0.1 | Penalizing large weights. |
| Precision Training Mode | bfloat16 | Utilizes bfloat16 for mixed precision training. |
| Optimizer | Adam | Optimization method for updating weights. |
| Loss Function | Cross-Entropy | Loss function for training the model. |

### 3.3 Protein Sequence Generation

For the distribution 𝒫 _Θ_(*S* |***P***) learned by the InstructPLM, we use nucleus sampling [47] and temperature scaling to generate sequences. We first employ a temperature parameter *τ* to modulate the sharpness of the distribution,

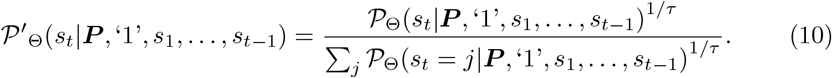

A lower temperature value causes the distribution to become sharper, reducing the diversity of designed protein sequences. Then, nucleus sampling is adopted for truncating the distribution to retain a subset (nucleus) that cumulatively covers a probability mass threshold specified by a hyperparameter top-p *p*. This method mitigates issues associated with other sampling approaches, such as the high randomness of pure sampling or the deterministic nature of greedy and beam search strategies. Formally, for the next token *s*_*t*_ over the vocabulary *V*, the smallest set *V* ^*′*^ of tokens is chosen by:

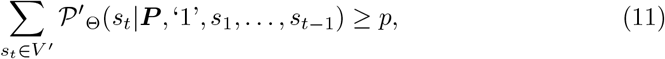

Finally, we sample new tokens from *V* ^*′*^ based on 𝒫 ^*′*^.

In the context of generating fixed-length protein sequences, we stop the generation process when the InstructPLM attains the pre-defined length. When InstructPLM predicts the C-terminal side token before meeting the pre-defined length, we modify the token probability by setting the probability of the C-terminal side to 0 and continuing the generation process.

In the recovery rate (Table 1), and DEDAL (Figure 2(A)) benchmark, we first reproduce the result of three baselines (ProteinMPNN, PiFld, ESM-IF) utilizing the official codebase with temperature *τ* = 0.15. We generate 20 sequences for each structure and report the mean recovery rate of 20 sequences as the final result. For InstructPLM, we adopt the fix-length generation for the recovery rate benchmark with the same temperature and a top-p of 0.9. In terms of the DEDAL benchmark, we let InstructPLM generate open-ended sequences with the same setting (*τ* = 0.15, *p* = 0.9) for fair comparison.

When designing functional proteins, for a comprehensive exploration of the sequence space, we scale up our generation process to produce 10,000 sequences with a temperature of 0.8 and a top-p of 0.9. This extensive generation process facilitates a robust evaluation of the InstructPLM’s ability to produce a wide array of plausible protein sequences conditioned on structural data, thereby enabling a more thorough assessment of the model’s performance and utility in computational biology and protein engineering tasks.

### 3.4 Experiment Details of Functional Proteins

We evaluate InstructPLM on two different proteins. PETase, a recently identified enzyme with the capability to degrade polyethylene terephthalate (PET), a common plastic, poses a test of InstructPLM’s ability to improve upon naturally occurring proteins. To further validate the generality and applicability of InstructPLM, we also investigate the designed sequences on L-MDH. L-MDH (Uniprot: A0A319AA41) is an important enzyme involved in the citric acid cycle, represents a fundamental metabolic process, and the absence of its definitive structure necessitates predictive modeling, testing the algorithm’s ability to function in the absence of solid experimental data. These proteins are chosen based on their varied structures and biological functions, presenting diverse challenges for InstructPLM and allowing us to probe the algorithm’s versatility and robustness.

#### 3.4.1 Enzyme Expression and Purification of PETase

The recombinant bacteria were inoculated into LB liquid medium (100 μg*/*mL ampicillin) and grown overnight at 37°C/200 r.p.m. for 14 h. The 2% cell cultures were transferred into 100 mL LB liquid medium (100 μg*/*mL ampicillin) and were shaken at 37 ^*°*^C /200 r.p.m. for 3 h. When the optical density at 600 nm (*OD*_600_) reached 0.6-0.8, isopropyl-1-thio-*β*-D-galactopyranoside (IPTG) was added to final concentrations of 0.1 mM and grown overnight at 20°C/200 r.p.m. for 24 h. Subsequently, supernatants were collected by centrifugation (8000 × g for 10 min at 4 ^*°*^C), and resuspended in 50 mM Tris-HCl buffer (pH 7.5) after washing twice with buffer (pH 7.5). The washed cells with 1 mg/mL lysozyme were disrupted by sonication (3-sec pulse on, 5-sec pulse off, AMP 35%). Cell debris was removed by centrifugation at 4 ^*°*^C, 8000 × g for 1 h, and the supernatants were filtrated by a 0.22 μm filter (Choice filter, Thermo Scientific). To obtain the purified enzymes, the supernatant, the samples were then applied to a 5 μg*/*mL HisTrap HP column (GE Healthcare). After washing unbound proteins (20 mM imidazole), the target protein was collected by 250 mM imidazole. The concentration of protein was determined using a protein assay kit (BCA Protein Assay Kit; Genstar, Beijing, China) with BSA as the standard and the purity of each protein was checked by SDS–PAGE analysis.

#### 3.4.2 PET Depolymerization Assay

To evaluate the variants’ activities, the amorphous gf-PET film (Goodfellow, 1 mm thickness, Ø 6 mm, roughly 67 mg) was soaked in 1950 mL of 50 mM Tris-HCl buffer (pH 8.5) and 50 μL of crude enzyme was added every 24 hours at of 40 ^*°*^C. The reactions were terminated by heat treatment (100 ^*°*^C, 10 min). The supernatant obtained by centrifugation (8,000 × g for 5 min) was then analyzed by high-performance liquid chromatography (HPLC, Agilent 1200 and Ultimate 3000 UHPLC systems), equipped with a Welch Ultimate XBC18 column (4.6 × 250 mm, 5 μm, Welch Materials, Inc., Shanghai, China) to quantify PET monomers released from PET depolymerization. The mobile phase was 40% methanol with 0.12% acetic acid (pH 2.5) at a flow rate of 0.7 mL/min, and the effluent was monitored at a wavelength of 240 nm.

#### 3.4.3 Enzyme Expression and Purification of L-MDH

The E. coli DH5*α* was used for plasmid construction, and E. coli BL21 (DE3) was applied for gene expression. The gene was synthesized into the pET28 expression plasmid by the XX company. All BL21 (DE3) strains containing expression plasmids were cultivated in Luria–Bertani (LB) liquid medium (5 g/L yeast extract, 10 g/L tryptone, 10 g/L NaCl). And were inoculated at 37 ^*°*^C with constant shaking at 220 rpm. Terrific Broth (TB) medium, 24 g/L yeast extract, 12 g/L peptone, 4 mL glycerol, 2.31 g/L KH2PO4, and 12.54 g/L K2HPO4, was the fermentation medium of the engineered strains. The medium was supplemented with 50 μg*/*mL kanamycin (kan) according to the screening markers carried by expression plasmids. Shake flask fermentation was performed as follows: the engineered strains constructed with different expression plasmids were grown in 5 mL of LB liquid medium overnight at 37 ^*°*^C. Then, 50 μL of the seed liquid was used to inoculate in 5 mL of TB medium. Cells were cultured at 37 ^*°*^C until *OD*_600_ reached up to 2-4, then Isopropyl *β*-D-1-thiogalactopyranoside (IPTG) was added to a final concentration of 0.3 mM. The fermentation was allowed to continue at 28 ^*°*^C for an additional 10 h.

#### 3.4.4 L-MDH Activity Assay

For L-MDH activity, 1 mL of fermentation broth was collected and centrifuged at 15000 × g for 3 min to obtain cell precipitation at 4 ^*°*^C. The cell precipitates were then resuspended in 1ml of phosphate-buffered saline (PBS). The cell suspensions were lysed by sonication on ice: 30% power, working 5 s, interval 5 s for 10 min, to obtain the MDH crude enzyme solution. The total reaction system is 200 μL, including 50 mM PBS (176 μL), 200 mM oxobutanedioic acid (2 μL), 200 mM NADH+, and 20 μL dilution of crude enzyme solution. Then the change of absorbance at 340 nm was detected and read absorbance data every 10s within 3 min. The enzyme activity was calculated according to the following formula: U*/*min = [Δ A/min] × [1/*ϵ*] × 1000 (*ϵ*= 6.22 L/(mmol/cm).

## 4 Related Work

Machine learning-based methods for capturing structure-sequence mappings have made significant progress in modeling protein sequences, but they still face challenges in achieving comprehensive and integrated modeling [48, 49]. To address the limitations of traditional approaches and investigate the long-range dependencies and dynamic interactions among protein residues, researchers have proposed various deep learning-based methods for protein inverse folding.

One prominent approach is to employ Graph Neural Networks (GNNs) to encode structural information and generate protein sequences. Models such as GraphTrans and StructGNN [30] encode structural information as node and edge embeddings, and iteratively decode the node embeddings into protein sequences using a conditional protein language model that captures both local dependencies and long-range interactions. Building upon this framework, GVP [32] introduces an additional scalar embedding based on dihedral angles to enhance the graph embedding. It follows a similar encoder-decoder pattern based on inter-residual message-passing and proposes a model quality assessment scheme for best-candidate selection. GCA [31] further improves the approach by proposing global-local context interaction modules, viewing the problem from the lens of language modeling to generate protein sequences through direct mappings. AlphaDesign [33] presents a simplified graph encoder and a constraint-aware decoder based on GVP [32]. By training with data from AlphaFoldDB [41], the framework captures effective constraints and accurate interactions among graph nodes. ProteinMPNN [6] capitalizes on the benefits of an auto-regressive encoding-decoding scheme and message-passing updating techniques. It enhances the structural position information of the Oxygen atom and pre-calculates **C**_*β*_ atom into protein structural representations, extending the computational design to multi-chain proteins and multi-scale noise adaptation. PiFold [34] improves the traditional encoding-decoding framework by introducing virtual atoms and backbone dihedrals, achieving equal-dimension protein encoders and a 70-fold acceleration compared to ProteinMPNN [6]. However, while effective in designing novel protein sequences, the diversity of generated sequences is limited by the small scale of training data.

To address the data limitation issue, ESM-IF [7] leverages the highly accurate protein folding of AlphaFold2 and trains a large-scale inverse folding framework based on GVP [32]. It utilizes approximately 16,000 high-quality protein sequence-structure data from CATH [35] and about 12 million data with noise obtained from AlphaFold2 [41], achieving high recovery of protein sequences but incurring a high computational cost. LM-Design [21] tackles the limited data problem by fine-tuning ESM models [17] and conducting inverse folding using effective embeddings from pre-trained protein models. It recovers design sequences through conditional mask predicting. However, the current approach does not promote natural protein sequences that exhibit dynamic lengths and differences due to its masked generation scheme.

In addition to methods that directly design sequences through structure, some approaches aim to expand the space of protein structures and achieve controllable generation by generating structurally similar distributions. RFDiffusion [50] utilizes a Transformer with symmetry equivariance, such as SE3 [51], as the denoising network of the Diffusion Model. By incorporating specific structural features or sequence information during sampling, the framework can generate a series of functional protein structures. Chroma [42] employs diffusion models as the framework and leverages equivariant networks along with additional structural classification information, substructures, shapes, and other conditions to achieve high designability and programmability. These methods provide new possibilities for protein sequence design through the generated structures, as the designed structures can be reverted back to sequence samples using inverse folding algorithms like ProteinMPNN.

Our framework applies sequence-to-sequence language models as autoregressive sequence decoders, which are more suitable for valuable sequence exploration compared to the masked generation scheme used in LM-Design [21].

## 5 Conclusion

We propose a novel protein design framework called InstructPLM that leverages cross-modality alignment and instruction fine-tuning techniques. This innovative approach enables the generation of proteins that are not only variable in length but also diverse, adhering to specified structural guidelines. Significantly, InstructPLM’s ability to produce variable-length sequences introduces a novel dimension to protein design – mimicking the natural evolutionary processes of insertion and deletion. This capability addresses a previously unmet need, as prior methods are limited to generating sequences of fixed lengths. Furthermore, the proteins designed by our framework, evidenced through rigorous experimentation, exhibit the desired functional characteristics, demonstrating the efficacy and utility of our approach. To this end, we believe that InstructPLM will serve as a powerful toolkit that facilitates advancements in drug discovery, therapeutic proteins, and beyond.

## 6 Appendix

### 6.1 Implementation Details

To ensure the reproducibility of our results and facilitate the understanding of our model’s architecture, we provide a detailed description of the hyperparameter and training settings employed in InstructPLM.

#### 6.1.1 Initialization

- **Linear Layers**: The weight parameters of the linear layers are initialized using a truncated normal distribution with a mean of 0 and a standard deviation of 0.02. The biases are initialized to zero.
- **Layer Normalization**: The weights and biases of the layer normalization modules are uniformly initialized to 1.0 and 0, respectively. This initialization scheme ensures a standardized starting point for the training process.

##### 6.1.2 Hyperparameters

We show our model hyperparameters and training configuration in Table 5.

##### 6.1.3 Tokenizer

We use the same tokenizer as the original ProGen2. The vocabulary of the tokenizer comprises 30 tokens, including 20 amino acids, 5 special tokens, and 5 unused tokens.

Among the 5 special tokens, we use ‘1’ and ‘2’ to represent the N-terminal and C-terminal sides of the protein sequence respectively, representing the end of the sequence by the end-of-sequence token (⟨*eos*⟩). To handle sequences with different lengths, the padding token (⟨*pad*⟩) will be added at the end of the sequences. Since the structure prompt is added at the beginning of the protein sequence to indicate the start of the sequence, thus we do not use the ⟨*bos*⟩ token. Fig. 5 shows an example of a constructed data batch.

##### 6.1.4 Training and Evaluation

During the training phase, following [6, 7, 21, 34] we first filter the CATH 4.2 dataset with sequence lengths less than 500. We exclude the loss calculation of the tokens where the corresponding structure is missing, as same as other special tokens: ’1’, ⟨*eos*⟩, and ⟨*pad*⟩. We keep the loss calculation of special token ’2’ to teach the model where to stop. We evaluate InstructPLM on the CATH 4.2 validation set at the end of each epoch, and perform early stopping based on validation LM-loss, which can drastically reduce the training cost. As the vocabulary of InstructPLM is larger than other baselines, for fair comparison, we only calculated the perplexity on 20 valid amino acids. The perplexity on the whole vocab is shown in Table 6. Our methods are still optimal even with a larger vocabulary. To evaluate designed sequences, we truncate each sequence between ‘1’ and ‘2’ from the original output of InstructPLM resulting in a sequence with pure amino acid tokens.

**Table 6.** Perplexity with/without special tokens.

| Model | Perplexity on CATH 4.2↓ |  |  |
| --- | --- | --- | --- |
|  | Short | Single-chain | All |
| InstructPLM-*full vocab.* | 3.28 | 3.20 | 2.71 |
| InstructPLM | 3.22 | 3.17 | 2.68 |

**Table 7.** Sequence design performance on TS50 and TS500. The metrics include perplexity (exponentiated categorical cross-entropy loss per residue) and recovery rate (percentage of correctly predicted residues). The best performance is **bold**, while the best baseline is indicated with an underline.

| Model | Perplexity ↓ |  | Recovery ↑ |  |
| --- | --- | --- | --- | --- |
|  | TS50 | TS500 | TS50 | TS500 |
| StructGNN [30] | 5.40 | 4.98 | 43.89 | 45.69 |
| GraphTrans [30] | 5.60 | 5.16 | 42.20 | 44.66 |
| GVP [32] | 4.71 | 4.20 | 44.14 | 49.14 |
| GCA [31] | 5.09 | 4.72 | 47.02 | 47.74 |
| AlphaDesign [33] | 5.25 | 4.93 | 48.36 | 49.23 |
| ProteinMPNN [6] | 3.93 | 3.53 | 54.43 | 58.08 |
| PiFold [34] | <u>3.86</u> | <u>3.44</u> | <u>58.72</u> | <u>60.42</u> |
| **InstructPLM** | **2.29** | **2.42** | **67.99** | **64.22** |
| (Relative Gain) | (-32.9%) | (-26.45%) | (+15.79%) | (+6.28%) |

### 6.2 More Results on TS50/TS500

We further test InstructPLM on the TS50 and TS500 datasets. As Table 7 shows, our methods are better than the best baselines by large margins.

### 6.3 Designed Sequences for PETase and L-MDH

As for PETase, Table 8 and Table 9 list the 15 designed protein sequences. Figure 6 shows the sequence alignment results by ENDscript [52] of PETase generated by InstructPLM, Fast-PETase[53] and wild-type PETase. As for L-MDH, Table 10 and Table 11 list the 15 designed protein sequences. Figure 8 shows the sequence alignment results by ENDscript [52] of active L-MDH generated by InstructPLM and wild-type L-MDH. An example of L-MDH designed by InstructPLM is illustrated in Figure 7.

**Table 8.**
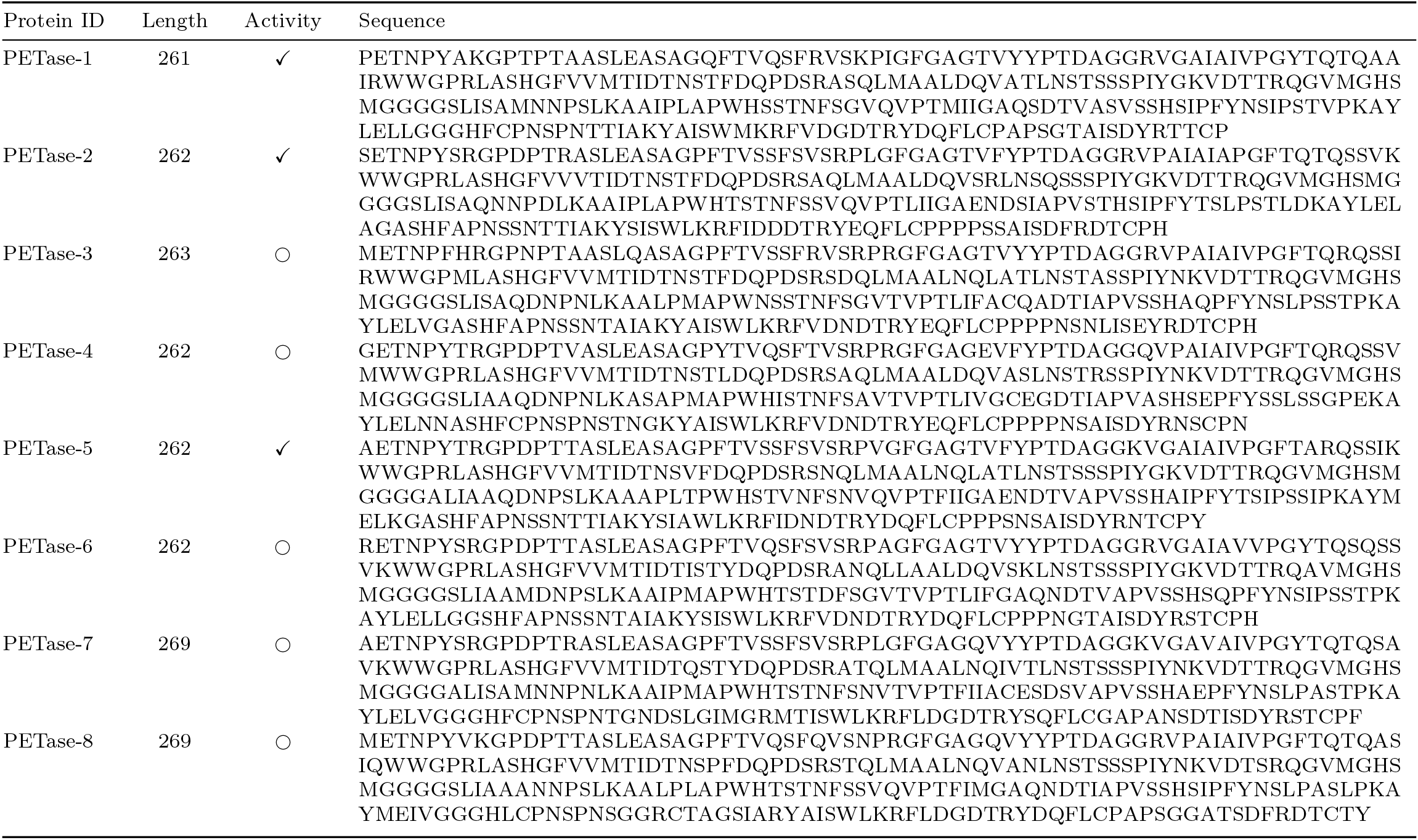
Designed PETase protein sequences (1-8), ✓indicates sequences where activity are detected; denotes sequences exhibiting activity superior to the wild-type.

**Table 9.**
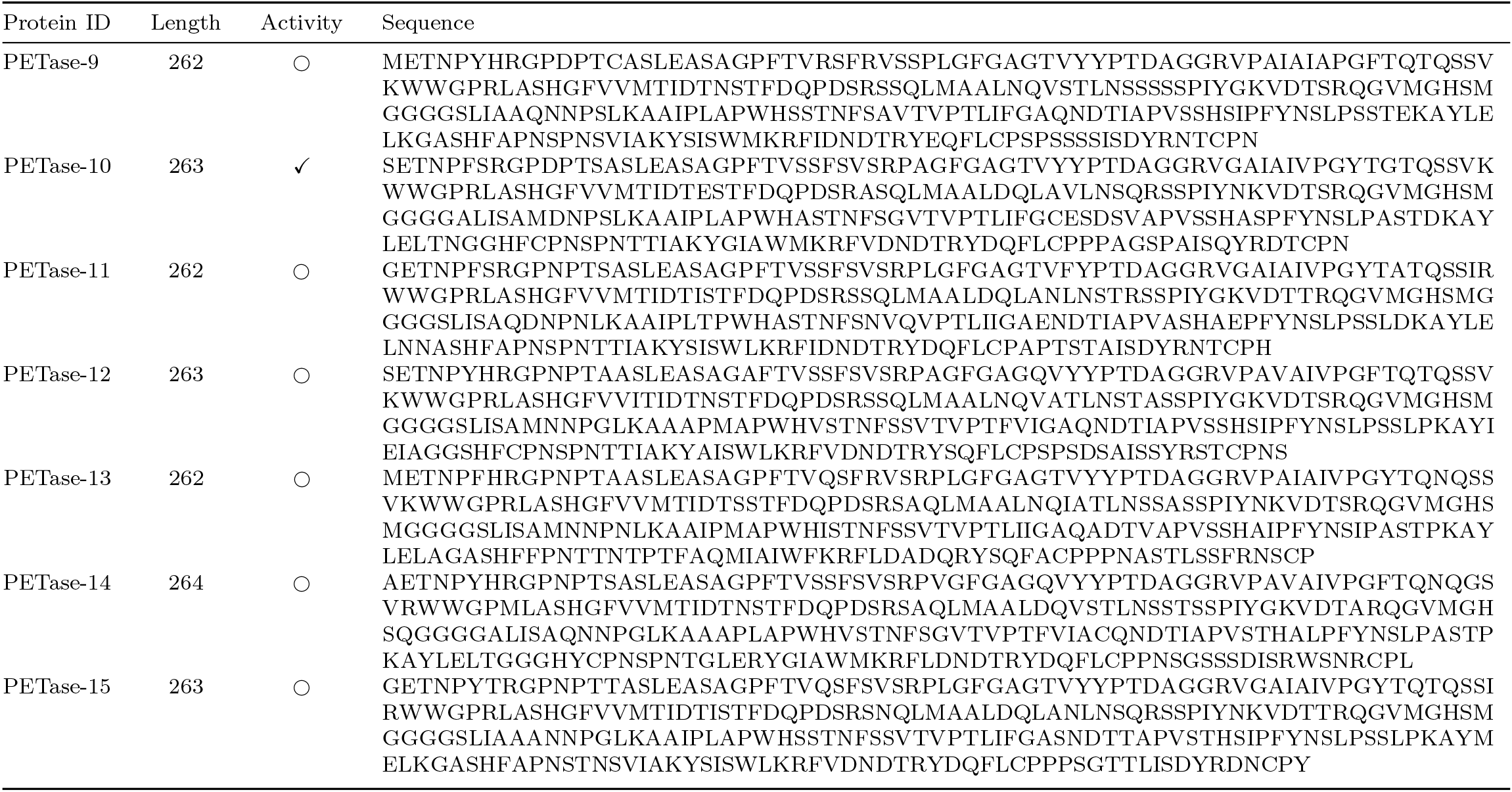
Designed PETase protein sequences (9-15), ✓indicates sequences where activity are detected; denotes sequences exhibiting activity superior to the wild-type.

**Table 10.**
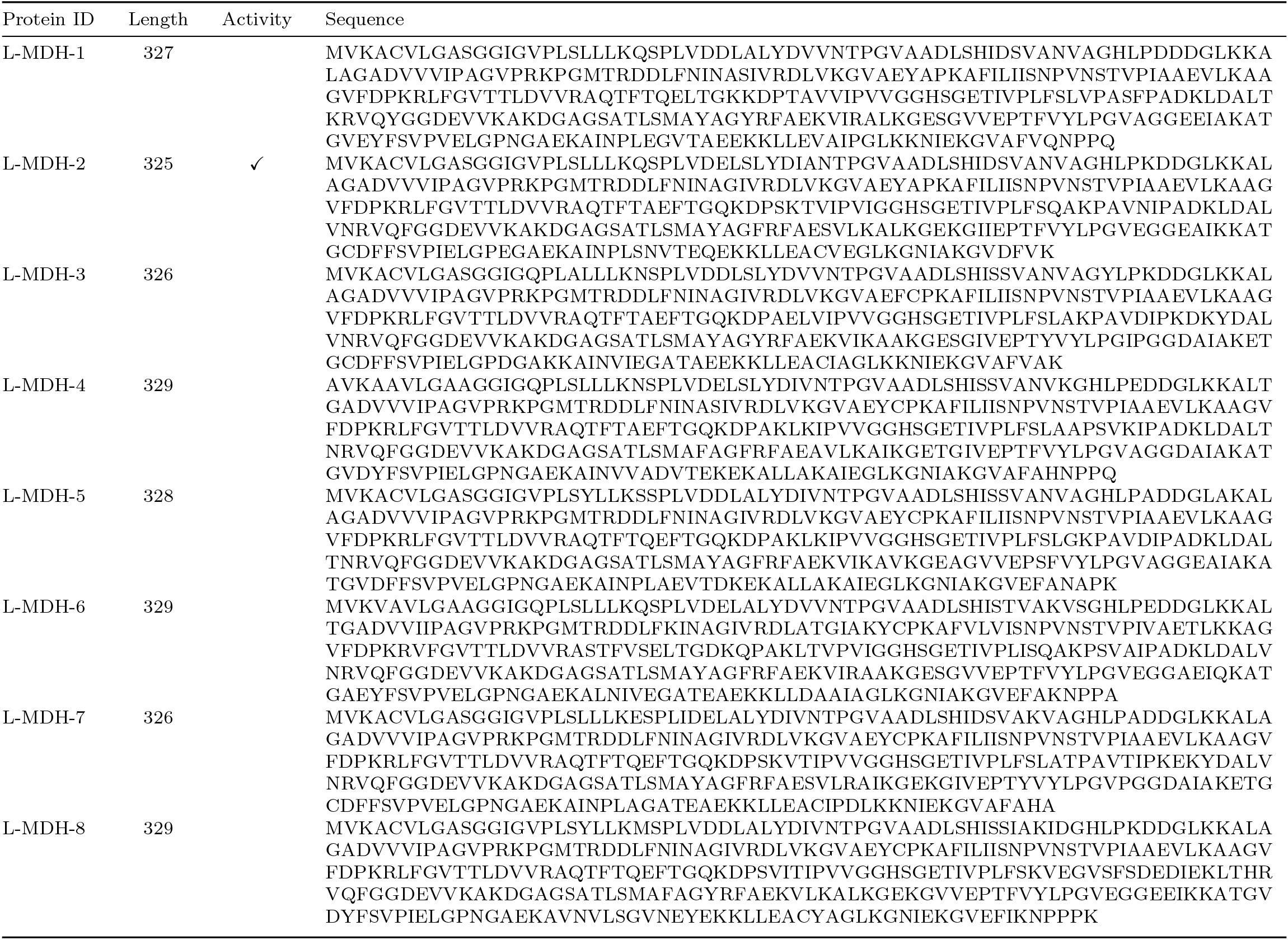
Designed L-MDH protein sequences (1-8), ✓indicates sequences where activity are detected; denotes sequences exhibiting activity superior to the wild-type.

**Table 11.**
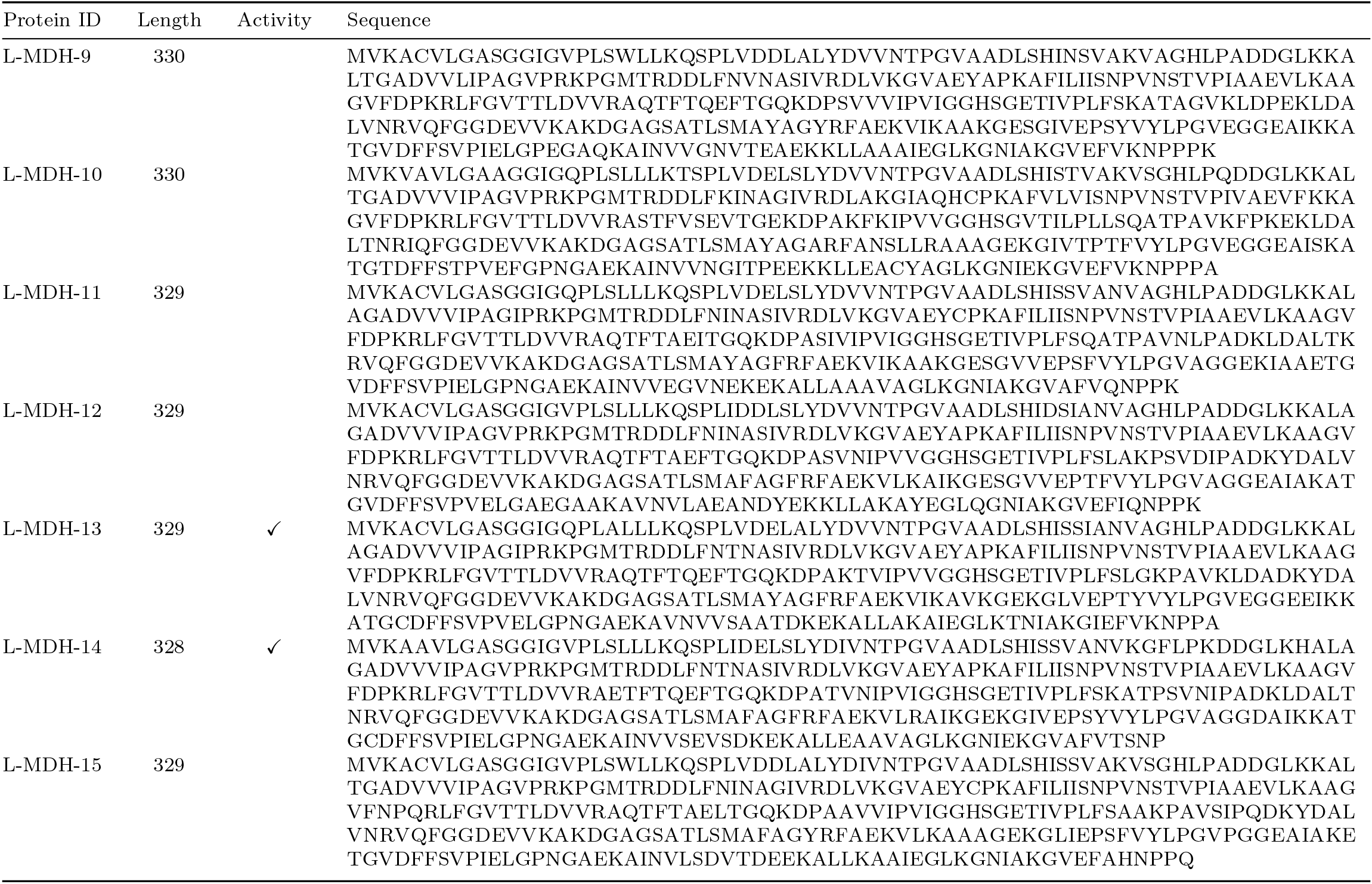
Designed L-MDH protein sequences (9-15), ✓indicates sequences where activity are detected; denotes sequences exhibiting activity superior to the wild-type.

## Footnotes

* Model weights: https://huggingface.co/InstructPLM/MPNN-ProGen2-xlarge-CATH42. Code: https://github.com/Eikor/InstructPLM.

1 https://github.com/dauparas/ProteinMPNN, including four vanilla models, two soluble models, and three *C*_*α*_ models.

